# Short hydrogen bonds enhance non-aromatic protein-related fluorescence

**DOI:** 10.1101/2020.01.22.915306

**Authors:** Amberley D. Stephens, Muhammad Nawaz Qaisrani, Michael T. Ruggiero, Gonzalo Diaz Miron, Uriel N. Morzan, Mariano C. González Lebrero, Saul T.E. Jones, Emiliano Poli, Andrew D. Bond, Philippa J. Woodhams, Elyse M. Kleist, Luca Grisanti, Ralph Gebauer, J. Axel Zeitler, Dan Credgington, Ali Hassanali, Gabriele S. Kaminski Schierle

## Abstract

Fluorescence in biological systems is usually associated with the presence of aromatic groups. Here, we show that specific hydrogen bonding networks can significantly affect fluorescence employing a combined experimental and computational approach. In particular, we reveal that the single amino acid L-glutamine, by undergoing a chemical transformation leading to the formation of a short hydrogen bond, displays optical properties that are significantly enhanced compared to L-glutamine itself. *Ab initio* molecular dynamics simulations highlight that these short hydrogen bonds prevent the appearance of a conical intersection between the excited and the ground states and thereby significantly decrease non-radiative transition probabilities. Our findings open the doors for the design of new photoactive materials with biophotonic applications.

**Significance statement:** Intrinsic fluorescence of non-aromatic amino acids is a puzzling phenomenon with an enormous potential for technological and biomedical applications. The physical origins of this effect, however, remain elusive. Herein, we demonstrate how specific hydrogen bond networks can modulate fluorescence. We highlight the key role played by short hydrogen bonds in the networks on the ensuing fluorescence and we provide a detailed molecular mechanism to explain this unusual non-aromatic optical properties. Our findings should benefit the design of novel optically active biomaterials for applications in biosensing and imaging.

## Introduction

Short peptides void of any aromatic residues have been shown to display an intrinsic fluorescence in the visible range (*1*, *2*). This has primarily been observed in fibrillar protein structures linked to neurodegenerative diseases, such as Alzheimer’s, Parkinson’s and Huntington’s diseases (*3*–*6*). Furthermore, optical properties of double amino acid based nanowires have also been reported, consisting either of two non-aromatic or two aromatic amino acids (*2*, *7*, *8*). We have previously suggested that the fluorescence of non-aromatic short crystal structures forming part of the amyloid-beta protein is enhanced by proton delocalisation (*5*). We have hypothesised that one of the prerequisites for this fluorescence observed in either amyloid structures or short peptide nanowires is related to hydrogen bonding or aromatic interlocks which, for the latter, decreases the bandgaps down to the semiconductive regions (*9*).

Despite our previous suggestion that proton delocalisation is strongly coupled to this intrinsic fluorescence, its direct role on putative fluorescing states has not been elucidated. We have thus searched for a model system, such as a single amino acid-based structure, that displays similar optical properties to amyloid fibrils and is permissive to more sophisticated computational approaches. We have been inspired by the small peptide nanostructures that have been pioneered by the Gazit laboratory (*9*) and by the fact that there are several neurodegenerative diseases that have been connected with an increased level of glutamines produced as part of a protein, as for example Huntingtin in Huntington’s disease which renders the protein more aggregation prone (*10*). It has been known that the amide group in L-glutamine (L-glu) is highly labile and thus can rapidly hydrolyse. We show here that the single amino acid L-glu upon heating in water can form a nanostructural material with optical properties similar to the ones observed in other amyloid fibrils such as in fibrils of amyloid-beta, alpha-synuclein or tau (*4*, *11*, *12*).

Using X-ray diffraction (XRD), we show that L-glu dissolved in water and upon heating becomes cyclysed forming a previously unreported structure which resembles L-pyroglutamine and which has been reported to be a component of amyloid-beta in the brain (*13*), but involves a low-barrier hydrogen bonded anionic dimer with an ammonium counterion. We have termed the new structure, i.e. L-pyroglutamine complexed with an ammonium ion, L-pyro-amm. L-pyro-amm has a microcrystalline plate morphology as shown by scanning electron microscopy (SEM). The newly formed solid has a unique hydrogen bond network formed by very strong hydrogen bonds (SHB) with a length of approximately 2.45 Å which is confirmed by using terahertz time-domain spectroscopy (THz-TDS).

By employing a combination of electronic structure calculations and *ab initio* molecular dynamics in the ground and excited state we provide an interpretation of the reported experiments. We illustrate the important contribution of different vibrational distortions on the optical properties. Furthermore, our simulations identify the origins of the non-aromatic fluorescence in L-pyro-amm, demonstrating the key role played by the SHBs, which prevent the appearance of a conical intersection, significantly reducing the chances of non-radiative relaxation towards the ground state and hence increasing the optical properties.

## Methods and Materials

### Experimental

#### Sample preparation of L-glutamine

L-glutamine (L-glu) (#G3126, #G8540, Sigma-Aldrich, Gillingham, UK) and L-pyroglutamine (L-pyro) (#83160, Sigma-Aldrich) were dissolved in 18.2Ω MilliQ H_2_O at a concentration of 0.3 M or 1 M. Aliquots were placed in a 65°C oven, since heating up proteins to 65°C increases the formation of amyloid structures as reported previously (*14*). Each aliquot was rotated to dissolve the powder once a day. Samples were either analysed in liquid form or dried on a glass or quartz cover slip (#043210.KG, Alfa Aesar, Lancashire, UK) either at room temperature (RT) or on a heat block set to 50°C. L-pyro-amm formed translucent crystals when dried.

#### Emission and excitation wavelength scans

Emission and excitation spectra were taken on a Hitachi F-4500 FL spectrophotometer (Hitachi High-Technologies Corporation, Tokyo, Japan) at room temperature (RT) in a quartz cuvette. For measurements, the excitation slit resolution was 5 nm or 10 nm and the emission slit resolution was 20 nm. The PMT voltage was set at 950 V and the scan speed set at 240 nm/min. The excitation scan was measured between 250 – 400 nm and the emission filter set to the emission maxima of the sample stated in the figure legend, with a slit resolution of 20 nm. The emission scan was measured between 380 - 560 nm and the excitation filter set to the excitation maxima of the sample stated in the figure legend, using a slit resolution of 5 nm. Four measurements were taken for each sample which were repeated at least three times and the background (air or H_2_O) was subtracted from the average.

#### Absorption measurements

Absorption measurements were taken on a UV-Vis-NIR Spectrophotometer, UV-3600 Plus (Shimadzu, Kyoto, Japan) and Cary 6000i (Agilent, Santa Clara, USA). 1 M or 0.3 M L-glu, L-pyro or L-pyro-amm solutions were measured in 10 mm QX cuvettes (Hellma Analytics, Müllheim, Germany) or dried on quartz coverslips. Measurements were taken between wavelengths 200 – 800 nm using 1 nm steps at a slow scan speed and a 1 nm resolution. The light source change wavelength was set at 393 nm and the grating change wavelength set at 750 nm. Both direct and integrating sphere measurements were taken and showed little difference in results, direct measurements are shown in the manuscript. Samples were measured at least three times and the experiments repeated at least three times, measurements were then averaged and H_2_O or coverslip only control was subtracted.

#### SEM (scanning electron microscopy)

SEM was performed using a FEI Magellan 400 HR-SEM at an acceleration voltage of 2 kV. L-pyro-amm samples were lyophilised by freezing in liquid nitrogen and freeze drying in a LyoQuest 85 (Telstar, Terrassa, Spain) and imaged on a glass coverslip.

#### X-ray diffraction

L-pyro-amm was dried on a glass coverslip in a 50°C oven and then at RT until crystals formed. Single crystal X-ray diffraction (SCXRD) measurements were performed at 180 °K with a Bruker D8-QUEST PHOTON-100 diffractometer, which utilised a Cu Kα radiation (λ = 1.54 Å), and an APEX-II CCD. Absorption corrections were made using SDABS, and data integration and reduction were performed with SAINT+. All non-hydrogen atoms were refined isotropically and anisotropically, followed by inclusion of the hydrogen atoms (determined using the excess electron density) and refinement isotropically. The structure is deposited in The Cambridge Crystallographic Data Centre, CCDC No. 1981551.

#### Terahertz Time-Domain Spectroscopy

All THz-TDS spectra were acquired using a commercial Terapulse 4000 spectrometer (TeraView Ltd, Cambridge, UK). Samples were prepared for THz-TDS measurements by diluting the solid air dried L-pyro-amm with polyethylene (~ 10% w/w concentration) by gentle mixing using an agate mortar, followed by pressing into 2 mm thick, 13 mm diameter pellets using a hydraulic press. All THz-TDS spectra shown are a result of division of sample and blank datasets, with the blank dataset represented the THz-TDS response of a pellet of pure polyethylene.

### Theoretical

#### Density functional theory (DFT)-THz Calculations

Calculations were performed using both the CRYSTAL17 (*15*) and Quantum Espresso (*16*) software packages. Geometry optimisations and vibrational analyses performed with the CRYSTAL17 code utilised the atom-centred split-valence triple-zeta 6-311g(2d,2p) basis set for all atom types. Based on a previous study related to ionic molecular crystals (*17*), the range-corrected WB97-X (*18*) functional was used. The vibrational analysis was performed within harmonic approximation, and infrared intensities were determined using the Berry Phase method (*19*). Energy convergence criteria were set to ΔE < 10^−8^ and 10^−11^ hartree for the geometry and frequency calculations, respectively.

#### Periodic time dependent (TD)-Density Functional Theory (DFT) Excited State Calculations

Simulations were performed using the fully periodic Quantum Espresso software package. The Becke-Lee-Yang-Parr (B3LYP) hybrid density functional was used with an energy cutoff of 40 Ry. The calculations of the excited state were performed within the framework of TDDFT using the Liouville-Lanczos formalism implemented in the freely available Quantum-Espresso package (*20*). In this approach, the optical spectra are obtained directly over the wide spectral range without taking into account the numerically complex calculations of the single exited states. We used plane wave basis set and the electron-ion interactions were taken into account via norm conserving Martins-Troullier pseudopotentials (*21*). To determine the ground state wave function, we used the gamma point of the Brillouin zone. All the periodic calculations employed the computationally demanding B3LYP (*22*) hybrid functional, the kinetic energy cutoff of 40 Ry was used for the wave functions. The intrinsic band width for the spectra was set to 0.003 Ry (~0.0408 eV).

#### Periodic Structure Geometry Optimisation

The structures obtained from the experiments were first geometrically optimized at 0°K using the Broyden-Fletcher-Goldfarb-Shanno (BFGS) minimisation algorithm implemented in CP2K (*23*, *24*) package. A convergence criterion for the wave function optimisation was used as 5×10^−7^ a.u.. Applying the method of the Gaussian and plane wave, the wave function was expended in the Gaussian double-zeta valence polarised (DZVP) basis set. The cutoff for the electronic density was set to 300 Ry. We used the gradient correction to the local density approximation and the core electrons were treated via Goedecker-Teter-Hutter pseudopotentials (*25*). In all the calculations, we used the Becke-Lee-Yang-Parr (BLYP) (*26*) functional with the D3(0) Grimme (*27*) dispersion corrections for the van der Waals interactions.

#### Ab Initio Molecular Dynamics Simulations

Ab initio Molecular Dynamics simulations (AIMD) were performed using Quickstep algorithm implemented in CP2K. In these calculations, the propagation of the nuclei was taken into account within the framework of the Born-Oppenheimer approximation. The simulations were performed in the NVT ensemble and the temperature was controlled during the simulations by using the velocity-rescaling thermostat (*28*). We used the time step of 0.5 femtosecond to update the nuclear coordinates and velocities while the total length of the simulations for each system is 50 picoseconds.

#### Excited State Cluster Calculations

A set of excited state calculations were performed on glutamine clusters in order to understand the role of the environment on the optical properties. Specifically, the optical properties of L-pyro-amm were investigated using various isolated cluster models with the Gaussian09 software package. The clusters were extracted directly from the crystal structure and used in various combinations (dimers, trimers, tetramers) to perform TD-DFT calculations. A split-valence triple-zeta 6-311g(2d,2p) basis set was used for all atom types together with the hybrid B3LYP functional. Some benchmark simulations, comparing the optical properties obtained from the periodic calculations using B3LYP to range corrected hybrid functionals like CAM-B3LYP, were also performed with these clusters.

We also performed a series of excited state optimisations on various model systems built from L-pyro-amm in order to examine the nature of the geometrical distortions that occur on the lowest electronic excited state. These calculations were also performed with the Gaussian09 software package. All clusters were surrounded with a continuum dielectric constant of 80, representing pure H_2_O. The 6-311G(2d,2p) basis set was used for all atoms together with the range corrected hybrid functional CAM-B3LYP (*29*). The clusters were first optimised in the ground state after which they were optimised on the first electronic excited state.

#### Non-adiabatic Decay Probabilities Using Excited State Molecular Dynamics

Excited state AIMD was employed, as implemented in the LIO quantum-chemical package (https://github.com/MALBECC/lio) (*30*–*33*), to analyse the influence of three key factors determining the optical properties of L-pyro-amm: (i) the formation of a SHB, (ii) the presence of ammonium, and (iii) the combination of ring deplanarisation and carbonyl stretching. Our model system for this study was the L-pyro-amm dimer with a single SHB. In order to assess the influence of the ammonium ion on the optical properties of L-pyro-amm, AIMD was performed both on the L-pyro-amm with and without the ammonium ion. Analogously, in order to shed light on the role of the SHB in the L-pyro-amm fluorescence, in addition to performing simulations of L-pyro at the natural SHB distance of 2.5 Å, we performed three replicas constraining the HB distance to 3.0 Å, 3.5 Å and 4.5 Å, respectively.

The initial structures for this analysis were extracted from the ground state AIMD described above. Subsequently, 3 ps ground state AIMD at 300K was performed to equilibrate the system, followed by 1ps of excited state AIMD. The non-radiative decay probability (NRP) was computed every time step using the TDDFT-based Trajectory Surface Hopping algorithm without permitting decays to the ground state (*34*–*36*). The non-adiabatic coupling elements between *S0* and *S1* (Equation 1) can be estimated analytically employing the method introduced by Tapavicza, *et al.,* (*37*).

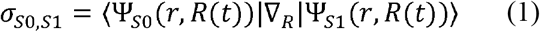

In this context, the non-radiative decay probability (NRP) from *S1* to *S0* (Equation 2) is expressed as:

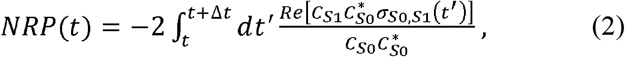

where C coefficients satisfy (Equation 3)

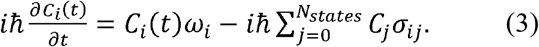

NRP(t) is the probability for the system to relax from *S1* to *S0* without fluorescent emission at time *t*. Hence, the accumulated non-radiative decay probability (Equation 4) accounts for the total relaxation probability from the excitation instant (*t=0*) up to time *t*.

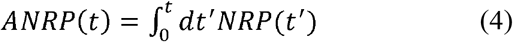

As mentioned above, this methodology was applied to five different initial conditions of the dimer at SHB, 3.0, 3.5, and 4.5 Å of hydrogen distance and for the dimer-amm at SHB distance. In order to avoid introducing large structural deformations, the distance constrains in the HB were introduced in combination with position restrains on the nitrogen and the C_β_ of the pyroglutamic ring.

The excited state AIMDs were performed employing the density functional theory (DFT) using the hybrid functional PBE0 level (*38*), with a 6-31G basis set and a time step of 0.5 fs. The computation of excitation energies, excited-state gradients, and non-adiabatic coupling vectors, the Linear Response TDDFT method was used within the Tamm-Dancoff approximation (*37*, *39*).

## Results and Discussion

It has long been known that poly-glutamine can form amyloid-like fibrillar structures *in vitro*. The more glutamine residues in the poly-glutamine polymer, the faster the aggregation propensity of the polypeptide chain. This led us to investigate whether L-glu on its own, under conditions which normally promote fibril formation, such as an increase in temperature (*14*), was able to form structures with similar optical properties, as recently observed for amyloid fibrils (*5*, *40*, *41*).

We first investigated the structure of L-pyro-amm, which formed after incubation of L-glu for 8 days at 65°C, using SEM and observed crystal structures shown in Fig. 1a. However, in order to investigate whether L-glu had indeed changed its crystal structure arrangement we performed XRD analysis of the resulting material. In Fig. 1b we show the crystal structure of the heated L-glu structure, which we termed L-pyro-amm, and the published crystal structures of L-glu and L-pyroglutamine (L-pyro) in Suppl. Fig. 1. Note, the L-pyro structure was analysed as it displayed structural similarities to the newly formed L-pyro-amm. Figures were obtained from geometry optimisations of the nuclear positions of the atoms using experimental densities. L-pyro-amm consists of 8 pyroglutamine groups and 4 ammonium ions (144 atoms) complexed within the crystal (see Fig. 1b). In contrast, as shown in Suppl. Fig. 1a, L-glu consists of 4 glutamine molecules (80 atoms) in the unit cell which form hydrogen bonds involving the termini and side chain. Furthermore, as shown in Suppl. Fig. 1b, L-pyro consists of 12 pyroglutamine molecules (192 atoms) in the unit cell forming hydrogen bonds involving the NH and COOH groups.

**Figure 1.**
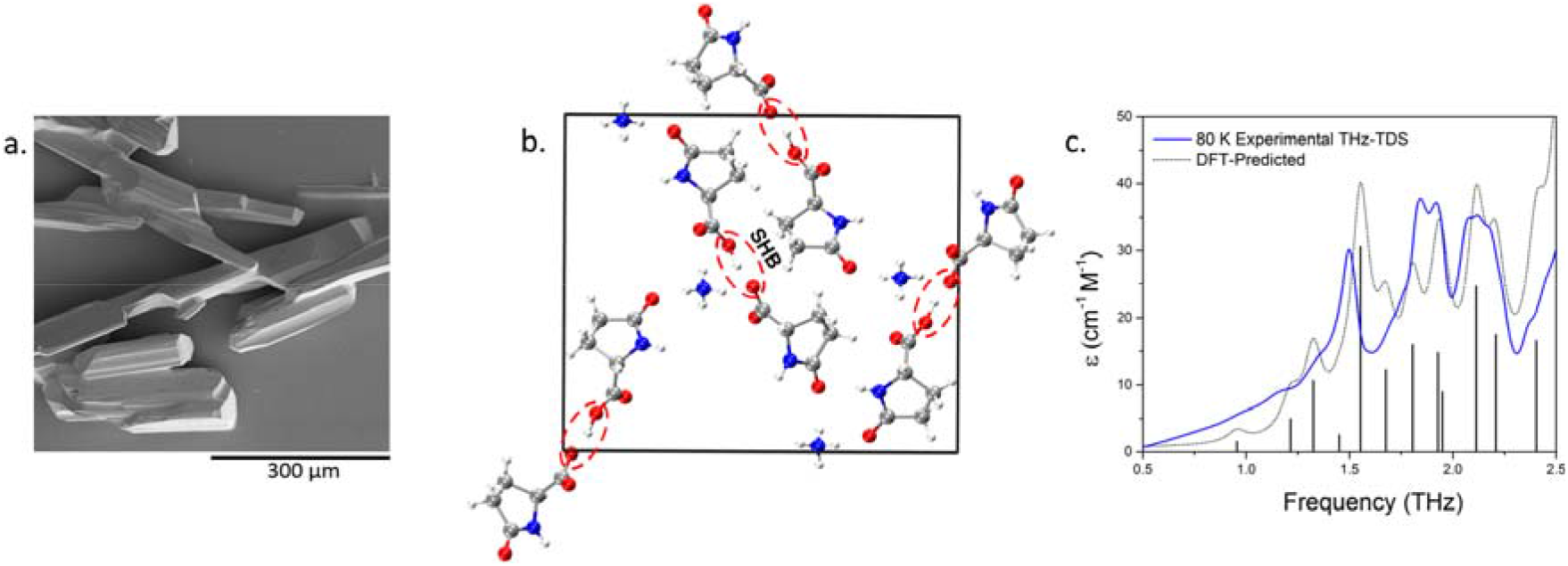
L-glu forms L-pyro-amm upon heating. (a) SEM image of crystals of L-pyro-amm dried. (b) XRD analysis of heated L-glu sample show the newly formed structure, L-pyro-amm. Geometry optimisations show that 8 pyroglutamine groups and four ammonium ions (144 atoms) are complexed in the crystal and a SHB of 2.45 Å (within red dashed lines) is present near the ammonium ion (white-hydrogen, red-oxygen, blue-nitrogen, grey-carbon). (c) Experimental (blue line) and theoretical (grey line) THz-TDS of the L-pyro-amm sample are in agreement and confirm the presence of the new L-pyro-amm structure.

L-pyro-amm has a rather unique hydrogen bond network structure since four of the pyroglutamine molecules are deprotonated and hence have a nominal negative charge, while the other four molecules are neutral. One of the important implications of this difference is that L-pyro-amm contains a very strong hydrogen bond. The red circled regions in Fig. 1b correspond to a short hydrogen bonds (SHBs) with a length of 2.45 Å, while those in L-glu and L-pyro range between ∼2.55-2.85 Å, Suppl. Fig. 1.

The structural change was further confirmed using THz-TDS measurements, as this technique is strongly dependent on the bulk packing arrangement as well as on the internal covalent structure of the molecules (*42*). The THz-TDS spectrum of the resulting solid, as well as the solid-state DFT predicted spectrum based on the single crystal XRD (SCXRD)-determined structure, is shown in Fig. 1c (full spectral assignment available in Suppl. Fig. 2). The agreement between the experimental and theoretical spectra further supports that full conversion of the sample occurs and thus enables additional investigations into the structural and electronic properties of the material. The agreement is also indicative of the ability of the theoretical model to not only reproduce the experimental structure, but also the weak forces found in solid structures.

We first investigated whether there were any differences in the optical properties associated with the three crystal structures. Comparing the absorption of L-glu, L-pyro and L-pyro-amm in water, we show that only L-pyro-amm has a significantly red-shifted absorption which lies in the 275-320 nm range, whereas both L-glu and L-pyro primarily absorb in the deep UV (<250 nm) (see Fig. 2a).

**Figure 2.**
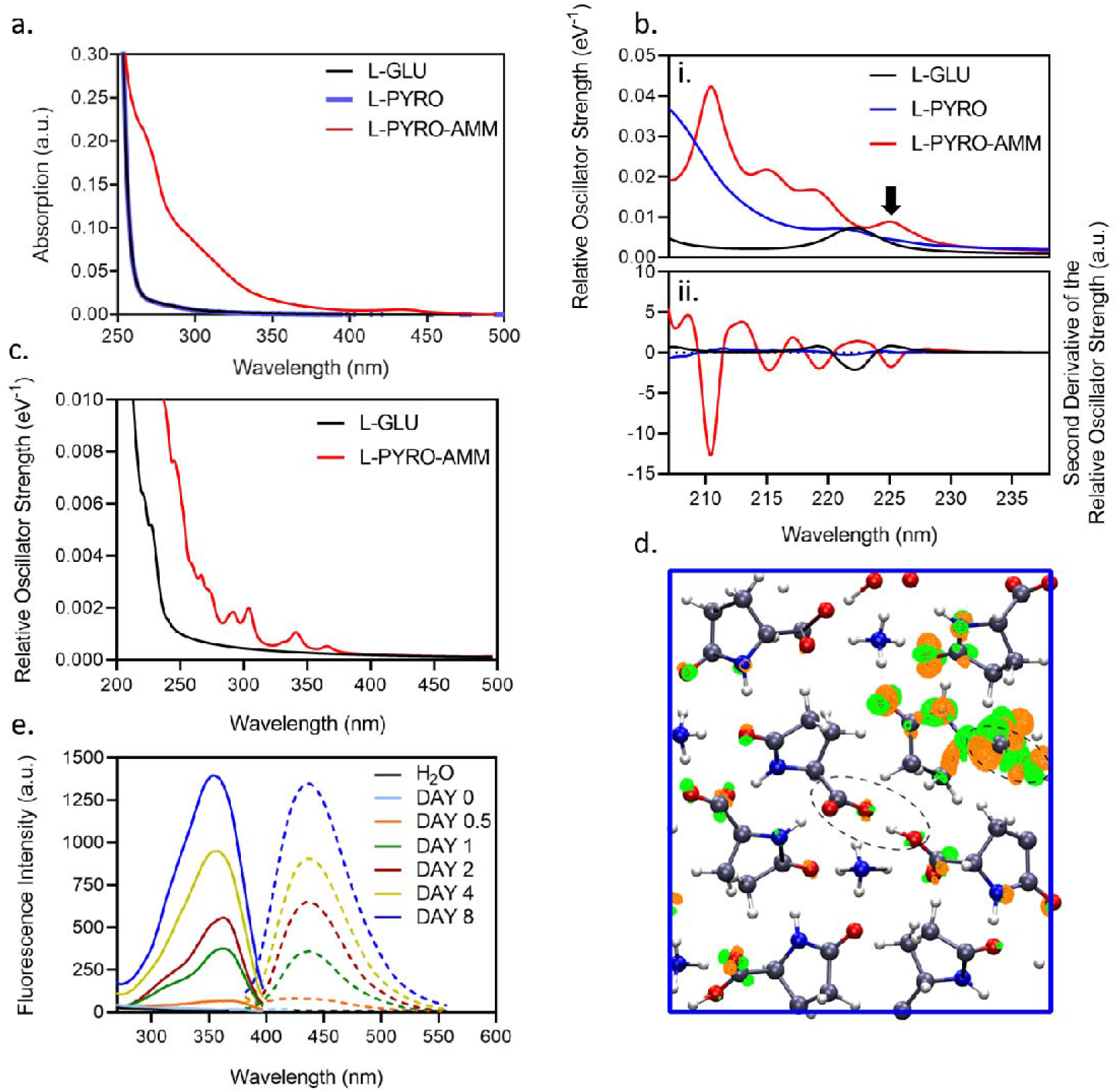
Optical properties of L-pyro-amm are distinct from L-glu and L-pyro. (a) Absorption spectra of 0.3 M L-glu (black), L-pyro (blue) and L-pyro-amm (red) (L-glu incubated for 8 days at 65°C) in water taken between 200 – 500 nm shows primarily features of L-pyro-amm. (b) Absorption spectra of L-glu, L-pyro and L-pyro-amm obtained from periodic density functional theory calculations with the B3LYP functional. L-pyro-amm features the lowest lying excited states which are characterised by the largest oscillator strengths. (c) Absorption spectra for L-glu and L-pyro-amm obtained from periodic simulations at room temperature. The spectra were computed by averaging over 25 frames randomly sampled from the *ab initio* molecular dynamics simulations. (d) The excited state electron density computed for L-pyro-amm from the optimised structure computed at the first peak (arrow in panel b). The lowest excited state density shows a response from various parts of the crystal structure including the pyroglutamic acid ring and the SHB region (see dashed circle). The orange and green surfaces correspond to regions involving a decrease and increase in electron density respectively, shown at an iso-value of 1×10^−5^. (e) 1 M L-glu was incubated at 65°C and the excitation and emission spectra were measured over time. Excitation spectra were measured between 250-400 nm with emission set at 420 nm and emission spectra were measured between 380-560 nm with the excitation set at 360 nm.

We next compared the experimental absorption spectra of L-glu, L-pyro and L-pyro-amm with the ones obtained from time dependent density functional theory (TDDFT). We highlight here, that the small size of the systems permitted us to determine the spectra using a hybrid functional, thereby not only advancing the quality of our theoretical predictions from previous studies (*5*, *40*, *41*) but also coupling the optical properties directly to different vibrational modes.

Fig. 2b illustrates the absorption spectra obtained for the TDDFT calculations on the 3 periodic systems in the ground state (i.e. at 0 °K). Panel b.i) shows the relative oscillator strength as a function of the frequency while panel b.ii) illustrates the second derivative of the oscillator strength permitting the positions of the maxima in the spectra to be more easily identified. The spectra reveal some striking differences between the different systems. Interestingly, we observe that L-pyro is essentially dark throughout the frequency range up to ~ 6eV. On the other hand, L-pyro-amm shows the presence of more structure in the spectrum. Specifically, it is the only system for which the spectrum features a low energy excitation at 226 nm (5.5 eV) and subsequently other peaks slightly above 220 nm (5.625eV) and 216 nm (5.75eV). While L-glu exhibits a peak at 222 nm (5.58eV), it is dark up to 206 nm (∼6eV).

We have previously shown that thermal fluctuations and in particular nuclear vibrations, such as proton transfer, have a large impact on the absorption spectra of peptide structures as compared to the 0 °K behavior (*5*, *43*–*45*). In Fig. 2c we show that, compared to the 0 °K spectra in Fig. 2b, thermal fluctuations cause a large red shift to around 3.4 eV (365 nm) for L-pyro-amm, close to what is observed experimentally. These spectra were computed averaging over 25 frames sampled from the molecular dynamics simulations. Interestingly, no such effect is observed for L-glu which remains weakly absorbing up to more than 5 eV (247 nm) as seen at 0 °K.

In order to understand better the physical origin of the low energy excitation at 226 nm (∼5.5 eV) in L-pyro-amm, we computed the electron response density at this frequency. This is illustrated in Fig. 2d, where we observe that most of the electron response is localised in regions around the pyroglutamine rings as well as regions near the SHB (see dashed circle in Fig. 2d). The optical response thus entails a charge reorganisation involving several parts of the molecular crystal. However, since L-pyro contains the same pyroglutamine rings as L-pyro-amm but absorbs more in the UV-spectrum, we conclude that the structural changes in the crystal in the presence of the SHB, which are the main features that distinguish L-pyro from L-pyro-amm, are responsible for the large Stokes shift observed in L-pyro experimentally and computationally.

We next investigated whether the above structures also display fluorescence excitation and emission properties as has been observed for amyloid-like structures reported previously (*5*, *40*, *41*). Fig. 2e shows the excitation scan from 250-400 nm (solid lines) with the emission set at 430 nm of L-glu in water at day 0 to 8 after incubation at 65°C. We observe an excitation peak at around 360 nm which is similar to what we have measured previously for amyloid proteins (*5*). The corresponding emission scan (dashed lines) with the excitation set at 360 nm and emission from 380-560 nm showed an emission peak around 430 nm, again lying in the same visible range as for amyloid fibrils. When the L-pyro-amm solution was dried the excitation and emission peaks were slightly blue shifted (Suppl. Fig. 3a) which may be due to a change in the molecular environment in the dried state. Importantly, we do not see any fluorescence in L-glu (without heating, i.e. at day 0 Fig. 2e.). To determine the importance of the ammonium ion experimentally, L-pyro was incubated in water and heated at 65°C for 8 days, and only a very weak fluorescence has been detected (Suppl. Fig. 3b).

As alluded to earlier, one of the factors that distinguishes L-pyro-amm from the other systems is the presence of the SHB (highlighted by red circles in Fig. 1b) and the presence of the ammonium ion. In order to characterise the behaviour of the SHB, we conducted *ab initio* molecular dynamics simulations of the three systems at 300 °K and examined the proton transfer coordinates defined as the difference in distance between the proton (H) and the two oxygen atoms (O1 and O2) that sandwich it and is commonly referred to as the proton transfer coordinate (d_O1-H_-d_O2-H_) as shown in Fig. 3 for different types of hydrogen bonds in the crystals. It is clear that the SHB in L-pyro-amm is characterised by a double-well potential. The barrier associated with this proton transfer is on the order of thermal energy, indicating that zero-point energy (ZPE) would make the proton transfer barrierless (*46*). An examination of similar proton transfer coordinates for hydrogen bonds in L-glu and L-pyro show that they are characterised by only single-well potentials.

**Figure 3:**
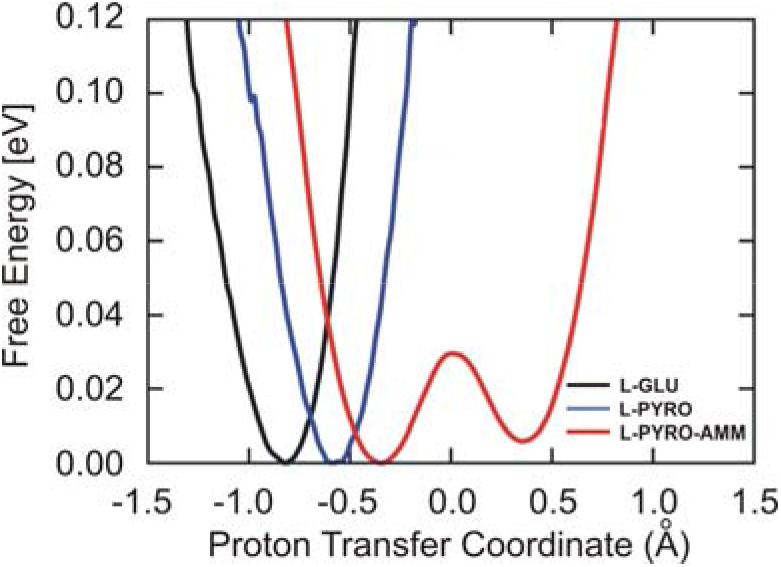
Free energy profiles along the proton transfer coordinate show only L-pyro-amm displays a double well potential. Free energy profiles along the proton transfer coordinate are displayed for L-glu (black), L-pyro (blue) and L-pyro-amm (red) at room temperature. The L-glu and L-pyro display single well potentials, while the L-pyro-amm system is the only one the exhibits a double well potential implying that there is proton transfer from one side to the other.

The nature of the optical properties is sensitive to the environment in which the glutamine molecules reside. It has previously been reported that charged amino acids already display an absorption in the range of 250-350 nm that is significantly red shifted (*47*, *48*). The origins of the low energy absorption were attributed to charge transfer excitations. The simulations of these systems were performed in the gas phase, rather than considering the protein environment such as shown for L-pyro-amm in Fig 2d. In comparison to the results presented in Fig. 2d, data presented in Fig. 4, show that the origins of the electronic transitions equally arise from a charge transfer (CT) between the highest occupied molecular orbital (HOMO) on the anionic dimer, and the lowest unoccupied molecular orbital (LUMO) centred on the ammonium cation when performed in the gas phase. Interestingly, the correct transition energy is only predicted when the ammonium cation is spatially near the centre of the dimer, which corresponds to the delocalisation of the negative charge and the SHB. Two generalised geometries, with the ammonium cation near the SHB (as seen in Fig. 4a) and away from the SHB (Fig. 4b), with the corresponding HOMO and LUMO orbitals are shown. The results predict a transition of 304 nm for dimer (a) and 669 nm for dimer (b), with dimer (a) most closely resembling the chemical environment present within the crystalline material.

**Figure 4.**
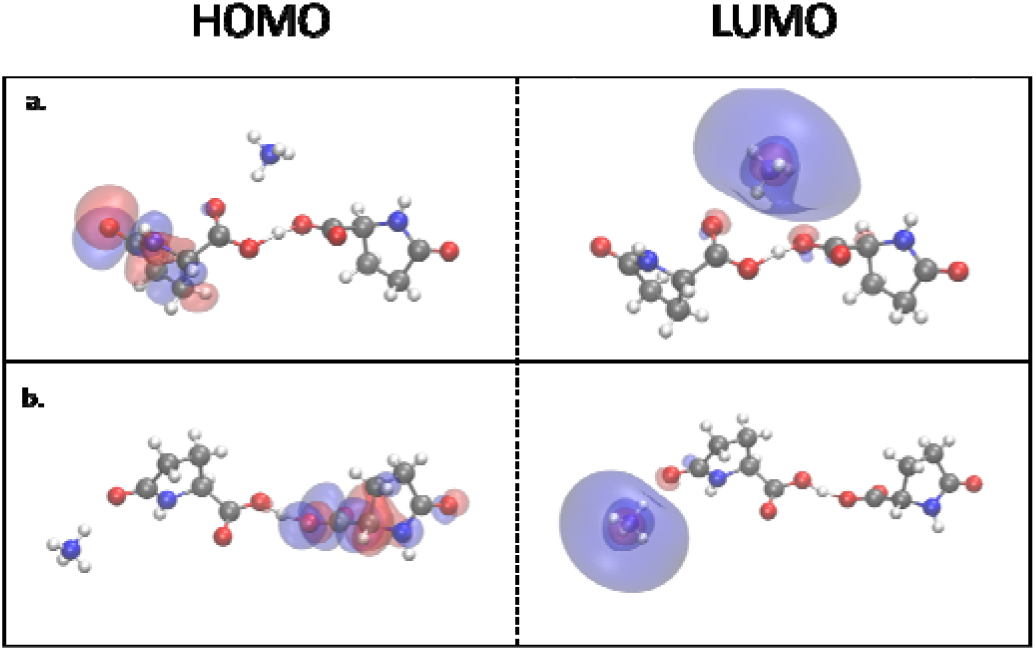
Comparison of the HOMO and LUMO orbitals on two L-pyro-amm models. L-pyro-amm structures are presented where the ammonium cation is located directly near the hydrogen bond (a), and where the ammonium cation is located away from the hydrogen bond (b). While both models predicted charge-transfer HOMO-LUMO states, only in the case of (a) is the transition predicted to be in the vicinity of the experimentally observed peak, 304 nm compared to 669 nm for (b).

The results show that charge transfer is capable to lead to absorption in the near UV when investigated in the gas phase, i.e. neglecting the direct protein environment. However, including the protein environment in the molecular crystal (Fig. 2d) results instead in the excitation being a charge reorganisation involving several different molecular groups of the crystal. Indeed, by shuffling the protons along the SHBs in the ground-state, we observe an electronic response involving the entire structural units of L-pyro-amm including both the hydrogen bonded regions and the pyroglutamic acid rings when the protein environment is accounted for (Suppl. Fig. 4).

Up to this point, we have shown that proton transfer along SHBs is an important part of the structural fluctuations in the ground state structure of L-pyro-amm. In order to characterise the nuclear relaxation upon photoexcitation, we conducted both excited state optimisations as well as excited state molecular dynamics simulations. We first show the results from the excited state optimisations obtained from clusters carved out from the different glutamine crystals and surrounded by a continuum dielectric constant of 80. Fig. 5a shows a scatter plot of the difference between the first excited state and ground-state energies as well as the corresponding oscillator strengths for the three systems, L-glu, L-pyro and L-pyro-amm. The scatter plots were obtained over the course of the excited state optimisation. We observe that the L-pyro-amm system is characterised by the largest oscillator strengths peaking at approximately 3.5eV (354 nm), which is consistent with our experimental findings. Although L-glu and L-pyro also have a peak at around 3.5 eV it is much weaker than the one of L-pyro-amm. We thus decided to focus on a series of excited state optimisations for various clusters of L-pyro-amm.

**Figure 5.**
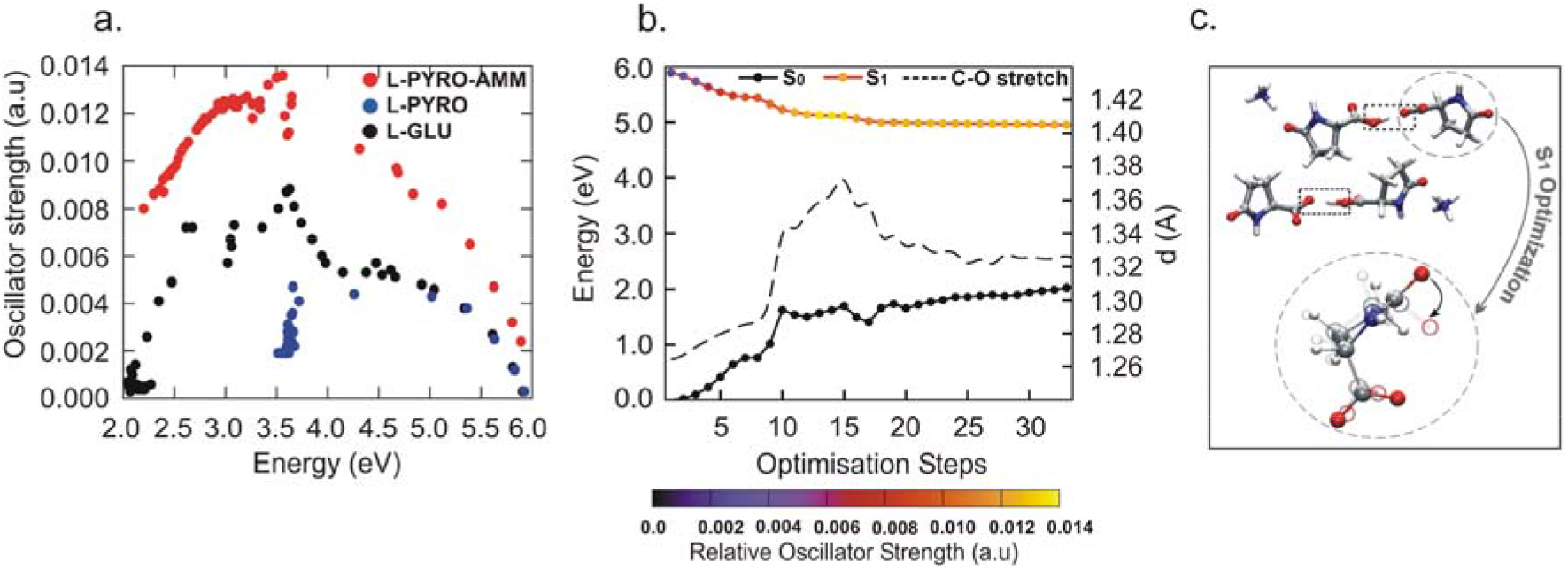
L-pyro-amm optical properties as seen through excited state optimisations. (a) Scatter plot of the oscillator strengths versus the emission energy (defined as the difference between the first excited state and the ground-state) during the excited optimisations of L-glu (black), L-pyro (blue) and L-pyro-amm (red). (b) Ground and excited state energies are plotted as a function of the excited state optimisation of the system shown in c. The curves on the excited state are colour-coded with the oscillator strengths. S_0_ and S_1_ refer to the ground and excited state energies, respectively. (c) A snapshot of the optimised excited state cluster also showing the lengthening of the carbonyl bond and the deplanarisation of the ring.

Fig. 5b shows the evolution of the first excited state energy for one of the L-pyro-amm clusters (Fig. 5c) across the optimisation steps. We find that the excited state energy drops by 0.5 eV combined with an enhancement of the oscillator strength while the ground state energy, shown in solid-black, rises by about 1.5eV. In this case, the closing of the energy gap to approximately 3eV is associated with an increase of the carbonyl oxygen bond as observed in previous studies (*48*–*50*), as well as a de-planarisation of the pyroglutamine ring (see Fig. 5c). Similar features are also observed in other clusters including one with the ammonium ion closer to the short hydrogen bond (Suppl. Fig. 5).

To further explore the preceding mechanisms, we next turn to deploying non-adiabatic decay estimations using excited state ab initio molecular dynamics simulations, as implemented in the LIO quantum-chemical package (https://github.com/MALBECC/lio) (*30*–*33*). This approach enables investigating the S0←S1 de-excitation probability providing a clear interpretation of the ensuing optical properties of L-pyro-amm (see Methods for details). The nature of the S0←S1 transition is mainly a ⍰’’=0 ← ⍰’=0 (where ⍰’ and ⍰’’ are the vibrational quantum numbers in the electronic ground and excited state respectively) with a FC factor between ground vibrational levels of ~0.82 (see Suppl. Fig. 6), therefore the excited state simulation was initiated by a vertical excitation to the S1 state. We performed two sets of excited state simulations: (i) varying the strength of the SHB to study its influence on the fluorescence of L-pyro dimer (Fig.6 a-b), and (ii) comparing L-pyro with L-pyro-amm dimers, to assess the role of ammonium ion on the transient excited state dynamics (Fig. 6c). Each trajectory was propagated for 1 ps from which the non-radiative decay probability (NRP) was determined by the fewest-switches trajectory surface hopping approach (*34*–*36*). Fig. 6d shows the time evolution of the accumulated NRP for various hydrogen bond lengths. The accumulated NRP (ANRP) represents the total probability for S_0_←S_1_ non-radiative relaxation. The larger value of the NRP implies that the system has a higher likelihood of decaying non-radiatively to the ground-state. On the other hand, a lower NRP would involve a longer excited-state lifetime increasing the fluorescence probability. Fig. 6d demonstrates unequivocally that the presence of the short hydrogen bond significantly reduces the chances of non-radiative relaxation towards the ground state. Examining the energy gap between the excited and ground state for this system shows that it occurs approximately ~3 eV (413 nm) (see Suppl. Fig. 7a) consistent with fluorescence in the blue/green visible regime. The presence of the ammonium ion further enhances, albeit by a subtle amount, the probability of trapping the system in the excited state.

**Figure 6.**
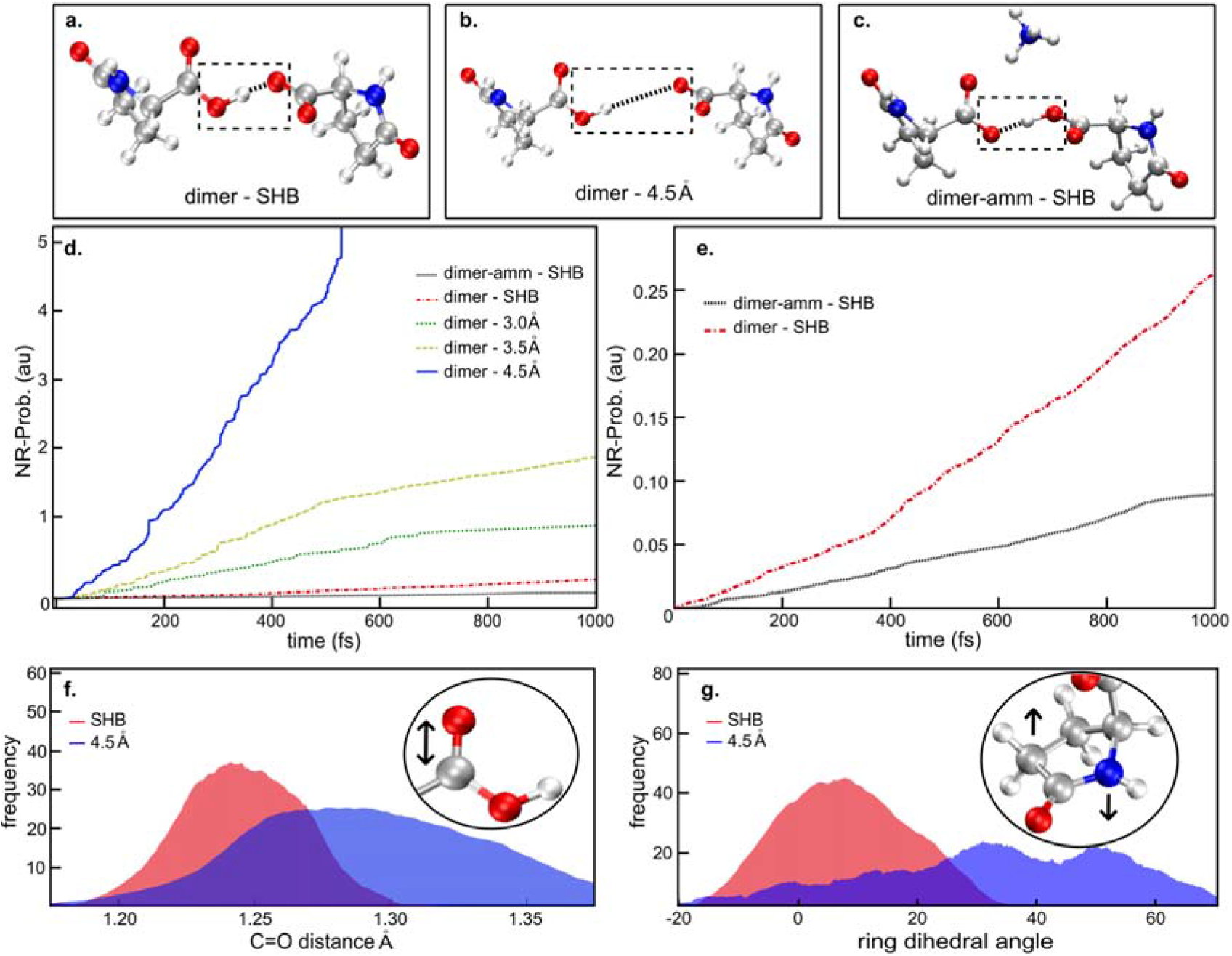
Excited state AIMD highlight how the presence of a SHB and the ammonium ion is enhancing fluorescence. Excited state AIMD performed for the (a) L-pyro dimer, (b) the L-pyro dimer constraining the SHB distance, and (c) the L-pyro-amm. (d) Accumulated non-radiative S_0_←S_1_ decay probability (ANRP) for L-pyro dimer (dimer-SHB) and the constrained SHB L-pyro (dimer-) where the SHB distance was fixed at 3.0 Å, 3.5 Å and 4.5 Å and for the L-pyro-amm dimer (dimer-amm-SHB). (e.) Accumulated non-radiative decay probability for L-pyro (red discontinuous line) and L-pyro-amm (black dashed line). (f) Carboxyl C=O distance histogram for the L-pyro-amm dimer at the SHB distance (red curve) or constraining the SHB bond to 4.5 Å (blue curve). (g) Ring dihedral angle histogram for the L-pyro-amm dimer at the SHB distance (red curve) or constraining the SHB bond to 4.5 Å (blue curve).

The excited state optimisations shown in Fig. 5, illustrate the important role played by the vibrational distortions in the excited state that on the ensuing optical property. In agreement with this, Figs. 6f and g show the distribution of the C=O bond length and the ring de-planarisation (computed as the sum of its internal dihedral angles) for the dimer systems with the SHB and weaker (longer) hydrogen bonds in the left and right panel, respectively. Interestingly, we see that the trajectories that undergo non-radiative decay are characterised by both, a significant increase in the carbonyl stretch (C=O bond length) by about a tenth of an Angstrom, and a large de-planarisation of the dihedral angle. The combined distortion of these vibrational coordinates leads to non-radiative decay as observed in the dimer system with longer hydrogen bonds. In the system with the short and strong hydrogen bonds, these modes are hampered which essentially prevents the system from easily accessing conical intersections. This increases the likelihood of observing fluorescence in the L-pyro-amm structure characterised by short-hydrogen-bonds. Therefore, both our experiments and simulations demonstrate the crucial role played by the SHB and in determining the fluorescence of L-pyro-amm.

## Conclusions

The experimental and theoretical findings presented here, elucidate a rather complex molecular mechanism associated with the non-aromatic intrinsic fluorescence in protein-like structures. In the case of L-glutamine, a chemical reaction creates a newly formed structure involving a cyclised pyroglutamic acid ring. This new structure features absorption in the UV and emission in the visible range very similar to the chemically distinct amyloid fibrils (*4*, *5*, *11*, *12*, *49*–*51*). The structural chromophore responsible for the optical properties in this new protein-related structure arises from a hydrogen bond network associated with structures involving short hydrogen bonds. Indeed, we have shown previously that similar to L-pyro-amm, the crystallised structure of the hydrophobic core of amyloid beta, 2Y3J, L-pyro-amm contains a SHB (*52*) leading to a double well potential in the ground state. The presence of strong hydrogen bonds along which proton transfer occurs and secondly, specific ionic interactions in close proximity, such as involving the ammonium ion of L-pyro-amm, affect the optical properties. Although the fluorescence observed in these systems is much weaker compared to conventional fluorophores, the physical and chemical properties of the hydrogen bond networks reported here maybe a generic feature across many other peptide structures.

Our non-adiabatic molecular dynamics simulations demonstrate that the presence of short hydrogen bonds with specific environmental conditions hinder vibrational deformations that can access conical intersections, as was previously speculated (*5*). The role of carbonyl groups is also consistent with a recent experimental study showing their importance for non-aromatic fluorescence (*53*). SHBs have recently been observed in different biological systems which have long been associated with either intrinsic fluorescence, such as NADP/NAD (*54*), FAD/FMN (*55*), the light-sensing chromophore in photoactive yellow protein (*56*), or in the active site of many enzymes, such as hydrolases and oxidoreductases (*57*, *58*), many of which consist of a highly complex H-bond structures similar to amyloid proteins. Furthermore, it has been recently reported that 1 in every 16 hydrogen bonds in over 1600 proteins are characterised by short hydrogen bonds (*58*). Thus, the mechanisms we espouse here may be more general. Our findings offer the possibility of designing novel biomaterial for applications in optical sensing or the design of novel biocompatible catalysts.

## Supporting Information

Supporting data include; Supplementary Figure 1. Clusters of L-glu and L-pyro. Supplementary Figure 2. Full spectral assignment of THz data. Supplementary Figure 3. L-pyro-amm has blue shifted fluorescence when dried and displays higher fluorescence intensity than L-pyro. Supplementary Figure 4. The optical properties of L-pyro-amm are sensitive to the environment and involves the electronic response of the entire structure. Supplementary Figure 5. L-pyro-amm undergoes vibrational distortions upon excitation as seen through excited state optimisations. Supplementary Figure 6. Potential energy surface along the SHB proton transfer coordinate in the ground (S0, panels A and D) and the excited (S1, panels B and C) states. Supplementary Figure 7. The energy gap between excited and ground state of L-pyro-amm is consistent with the experiments and eventually deactivates via a conical intersection back to the ground state. SCXRD data are available at the Cambridge Crystallographic Data Centre, CCDC No. 1981551. Raw data are available at the University of Cambridge Repository https://doi.org/10.17863/CAM.57945.

## Author Information

### Corresponding Authors

Ali A. Hassanali

Gabriele S. Kaminski Schierle

### Author Contributions

G.S.K.S and A.H conceptualised the manuscript. *A.D.S and M.N.Q contributed equally. A.D.S prepared samples for all experimental data. P.J.W. performed SEM experiments. A.D.B and M.T.R performed XRD measurements. M.T.R. performed THz experiments and DFT-THz calculations. A.D.S. performed excitation and emission measurements. A.D.S and S.T.J. performed absorption measurements. M.T.R and E.M.K performed TD-DFT cluster calculations. M.N.Q, E.P, L.G, R.G and A.H performed AIMD, Periodic TD-DFT Excited State Calculations and Periodic Structure Geometry Optimisation calculations. G.D.M, U.N.M, and M.C.GL performed excited state molecular dynamics. A.D.S, M.N.Q, M.T.R, S.T.J, L.G, G.D.M, U.N.M, M.C.G.L, J.A.Z, D.C, A.H and G.S.K.S contributed to manuscript writing. All authors have given approval to the final version of the manuscript.

### Notes

The authors declare no competing financial interest.

## Acknowledgments

We thank Bluebell Drummond, Talia Shmool and Chetan Poudel for work that is not in the final manuscript.

## Funding Sources

G.S.KS. acknowledges funding from the Wellcome Trust (065807/Z/01/Z) (203249/Z/16/Z), the UK Medical Research Council (MRC) (MR/K02292X/1), Alzheimer Research UK (ARUK) (ARUK-PG013-14), Michael J Fox Foundation (16238) and Infinitus China Ltd. M.T.R and J.A.Z acknowledge EPSRC funding (EP/N022769/1). P.J.W acknowledges EPSRC funding (EP/L016087/1). A.D.S. acknowledges Alzheimer Research UK the British Biophysical Society (BSS) for travel grants.

## Abbreviations

SEM: scanning electron microscopy
XRD: X-ray diffraction
THz-TDS: terahertz time domain spectroscopy
DFT: density functional theory
TD-DFT: time dependent DFT
SHB: short hydrogen bond
CT: charge transfer
L-glu: L-glutamine
L-pyro: L-pyroglutamine
L-pyro-amm: L-pyroglutamine complexed with an ammonium ion.

## Supplementary Information for

**Supplementary Figure 1.**
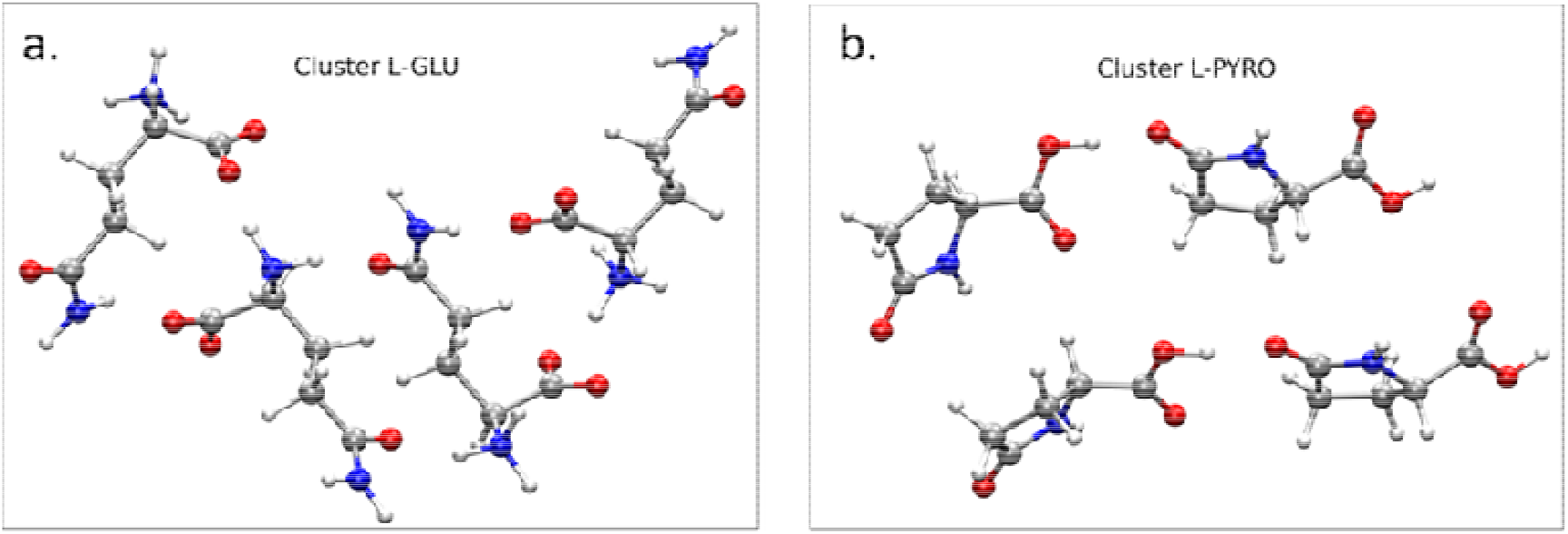
Clusters of L-glu and L-pyro. Structural representations of (a) L-glutamine (L-glu) and (b) L-pyroglutamine (L-pyro) clusters from the published crystal structure.

**Supplementary Figure 2.**
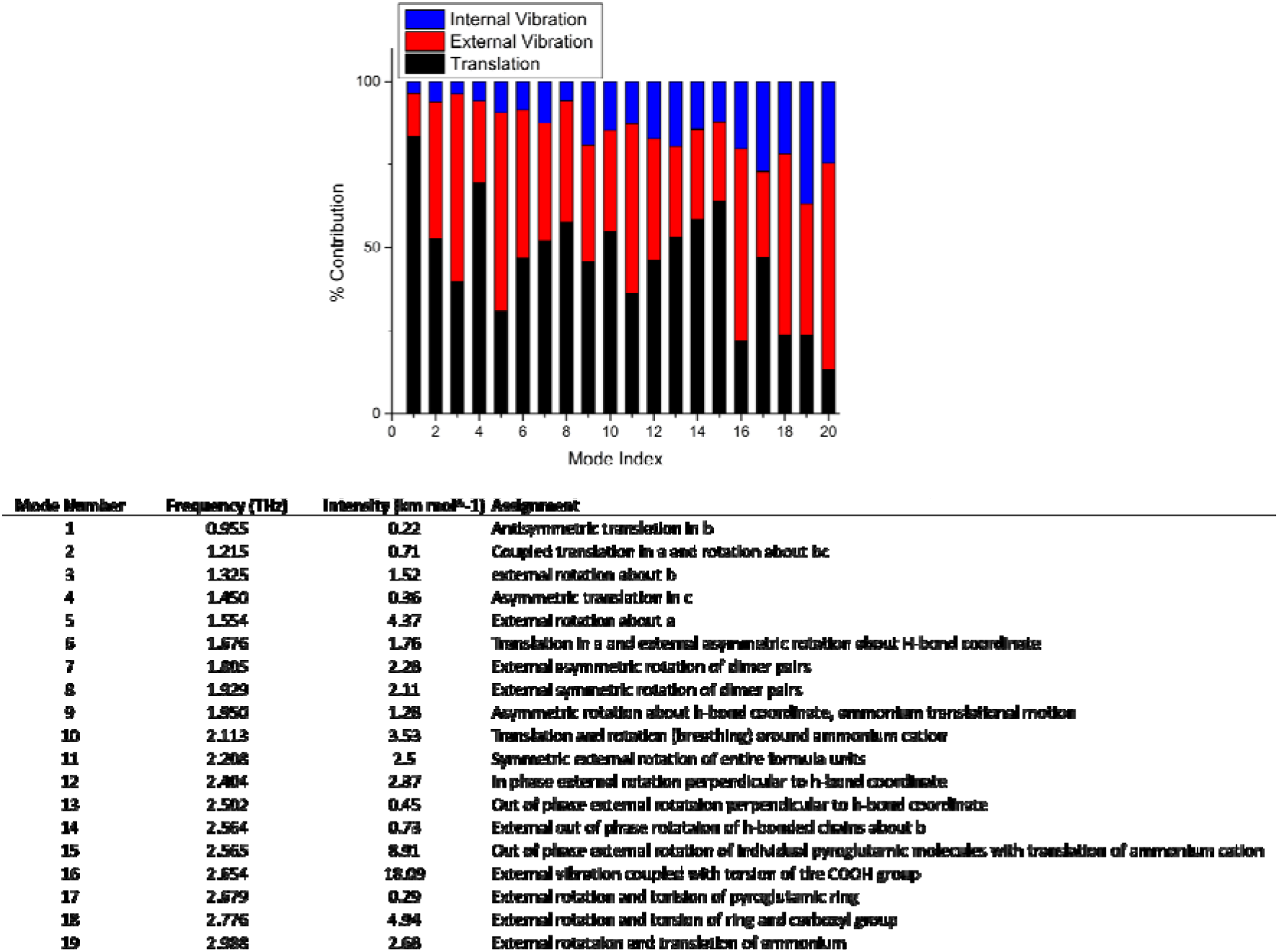
Full spectral assignment of THz data. The top chart shows the contribution to each IR-active mode including external translations and hindered rotations, and internal vibrational motions (i.e. torsions), while the bottom table lists the detailed assignment for each mode.

**Supplementary Figure 3.**
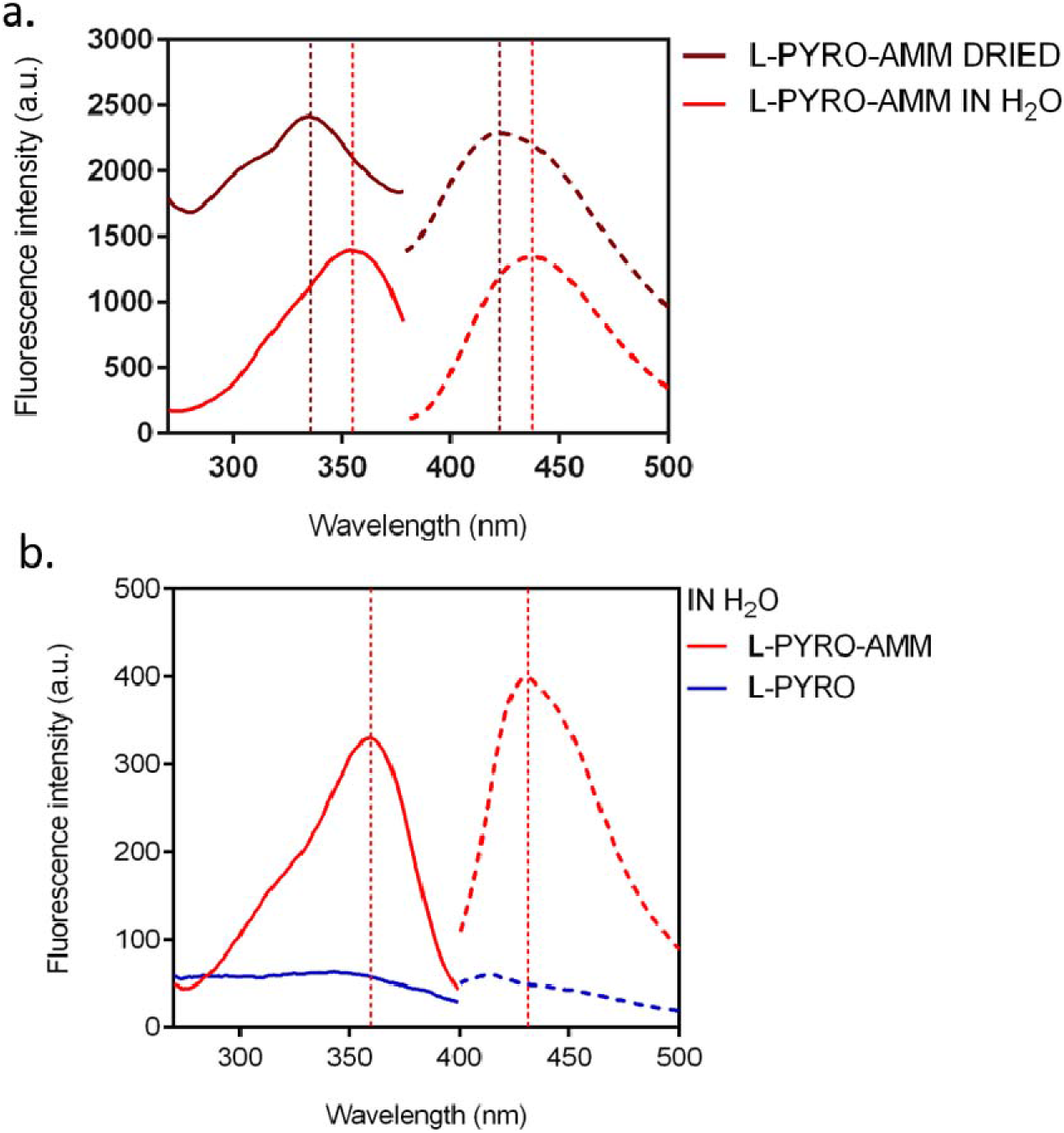
L-pyro-amm has blue shifted fluorescence when dried and displays higher fluorescence intensity than L-pyro. (a)The excitation peak of L-pyro-amm when dried (solid dark red line) is blue shifted with a peak maximum ~ 340 nm compared to L-pyro-amm in H_2_O (solid red line) which has a peak maximum ~ 360 nm. The emission peak of L-pyro-amm when dried (dashed dark red line) is also blue shifted, with a peak maximum ~ 420 nm, compared to L-pyro-amm in H_2_O (dashed red line) with a peak maximum ~ 430 nm. (b) 1 M L-glu and 1 M L-pyro (blue) were incubated in H_2_O for 8 days at 65°C. After 9 days, the L-glu had converted into the L-pyro-amm structure (red). L-pyro-amm has a clear excitation peak maximum at ~360 nm and emission peak maximum at ~ 430 nm, while L-pyro (blue), although not completely dark, has no clear excitation or emission peak.

**Supplementary Figure 4.**
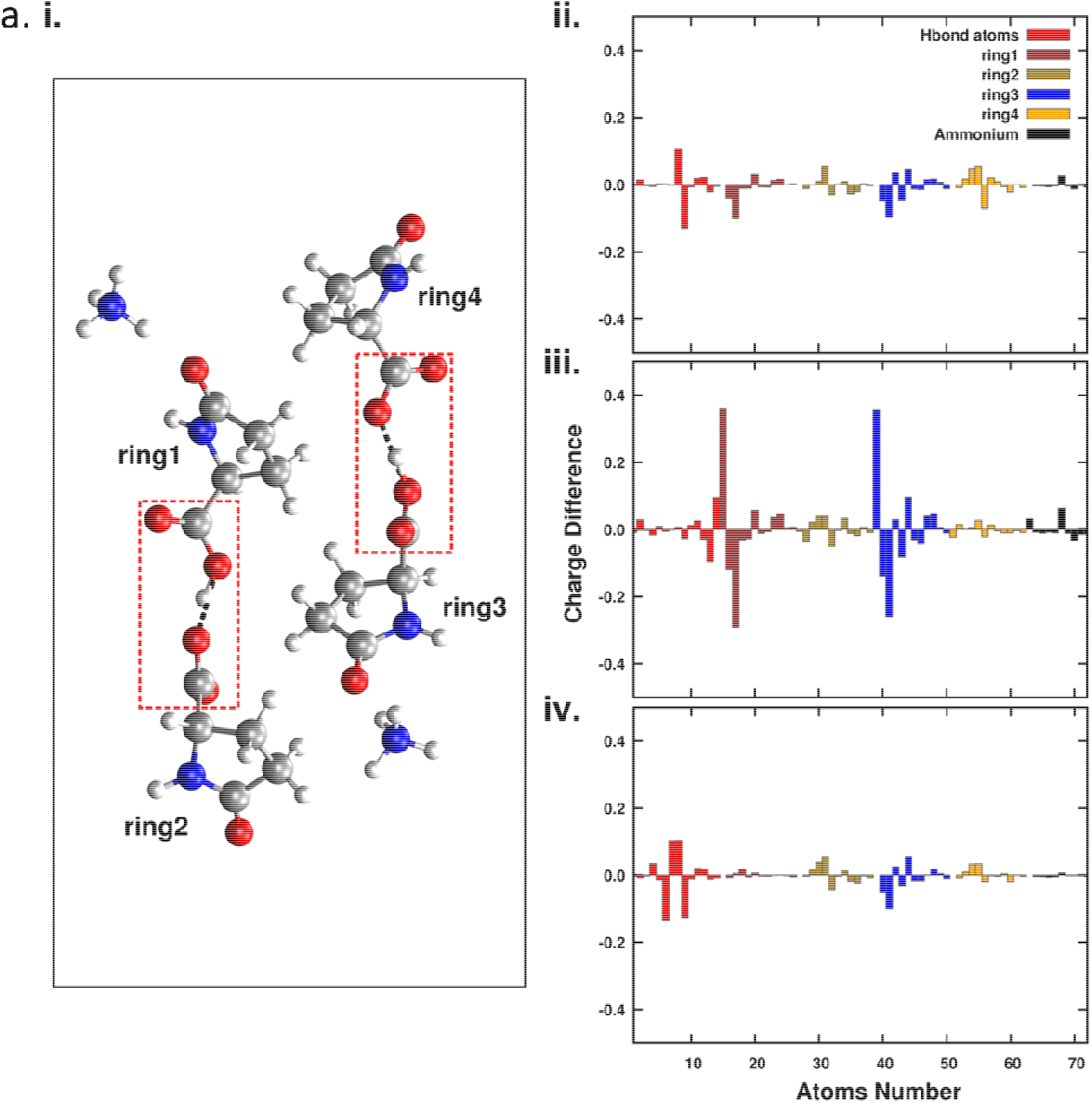
The optical properties of L-pyro-amm are sensitive to the environment and involves the electronic response of the entire structure. (a)(i) L-pyro-amm cluster used to examine the sensitivity of the optical response on different parts of the cluster upon moving different protons. (ii) Charge differences between the ground and excited state are computed using restrained electrostatic potential atomic partial charges (RESP) for the cluster shown in i). Note, the electronic response involves all the atoms of the cluster. (iii) The two protons in the rectangle regions are displaced to be in the centre of the hydrogen bond and the charge differences are then computed. As illustrated, the proton displacement leads to a larger change in magnitude of the charges. (iv) The charge differences are computed for another nuclear configuration for which the protons are kept fixed but the O—O distance is increased from 2.45 to 3.2 Angstroms. The charge differences obtained here are quite similar to the original condition shown in (ii).

**Supplementary Figure 5.**
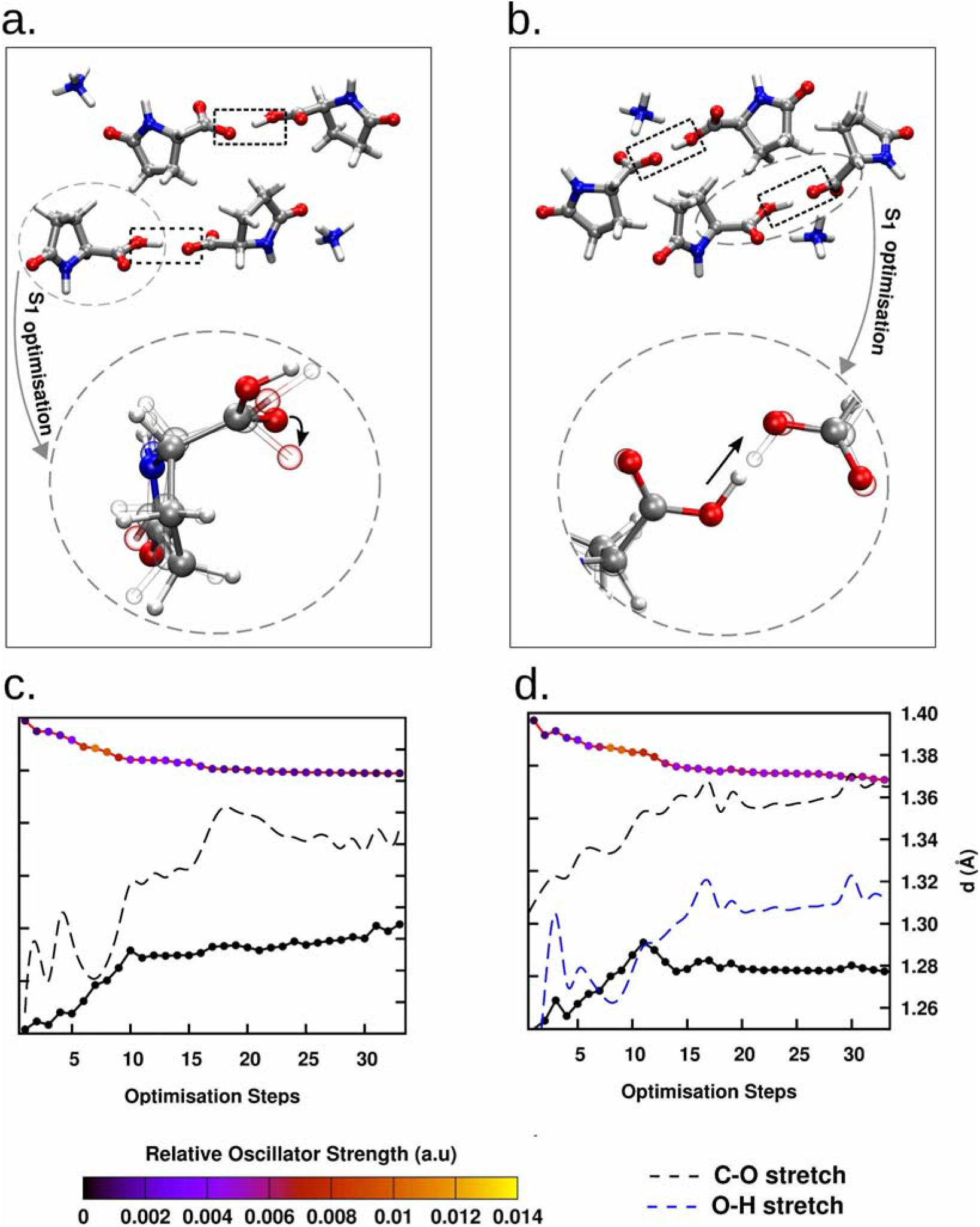
L-pyro-amm undergoes vibrational distortions upon excitation as seen through excited state optimisations. (a-b) correspond to two additional clusters to the one reported in the main text that were built from the L-pyro-mm crystal structure. c) and (d) correspond to the evolution of the excited state energy and ground-state energy over the course of the optimisation. Also shown, are the changes in the C-O bond lengths (dashed black) which increases on the excited state surface. In (d) we show the O-H bond stretch where a proton transfer is observed on the excited state.

**Supplementary Figure 6.**
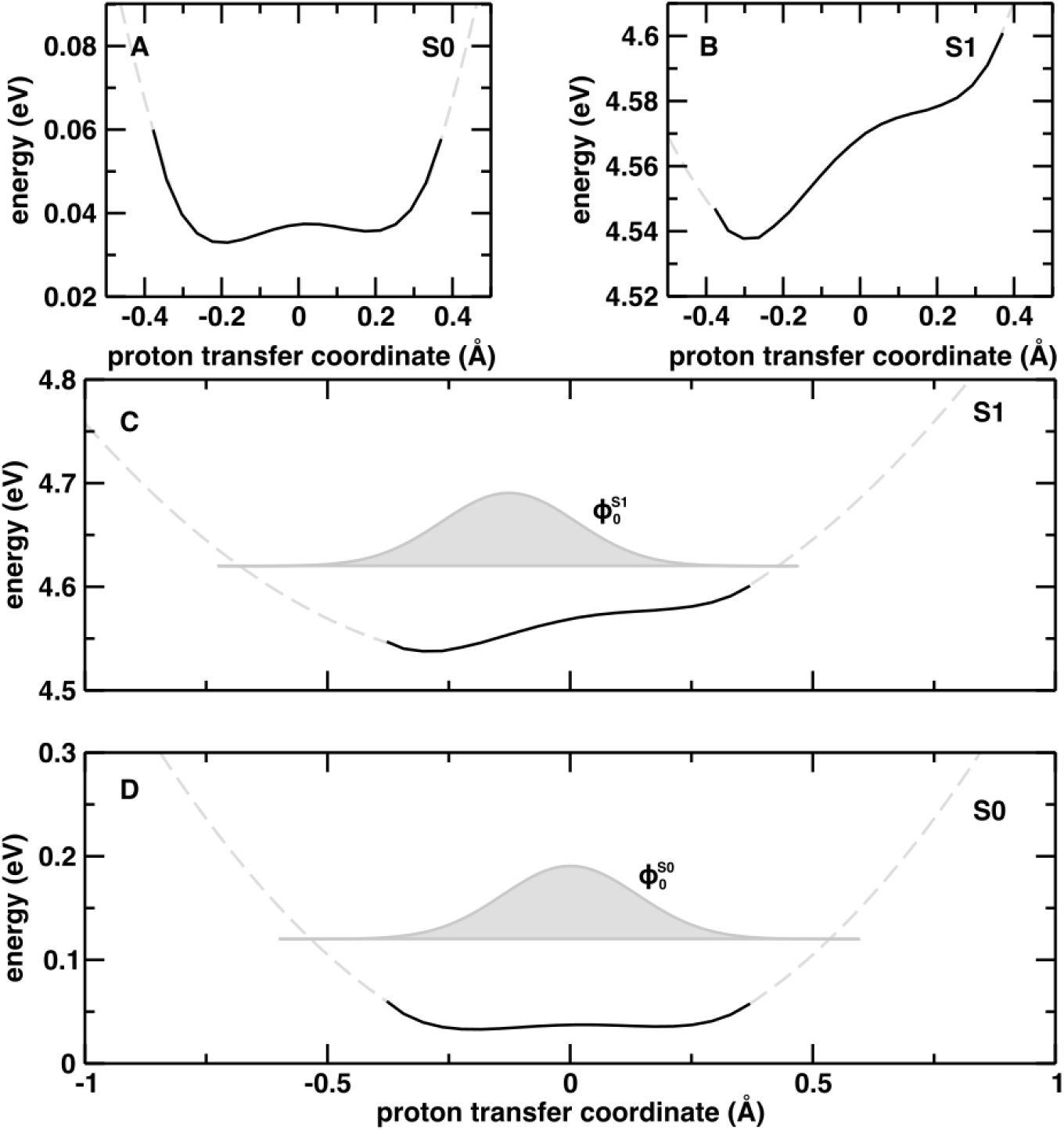
The S0 and S1 nuclear wavefunctions associated to the SHB proton transfer coordinate reveal a vibronic transition dominated by a 0←0 character. The calculated potential energy surfaces (black continuous lines) are prolongated with a quadratic fitting (gray dashed lines). Panels C and D illustrate the ground-state vibrational wavefunction in the S0 and the S1 electronic states (and respectively). The resulting Franck-Condon factor between and is 0.82, showing that the nature of the S1 ← S0 transition is mainly a ⍰’’=0 ← ⍰’=0 (where ⍰’ and ⍰’’ are the vibrational quantum numbers in the electronic ground and excited state respectively).

**Supplementary Figure 7.**
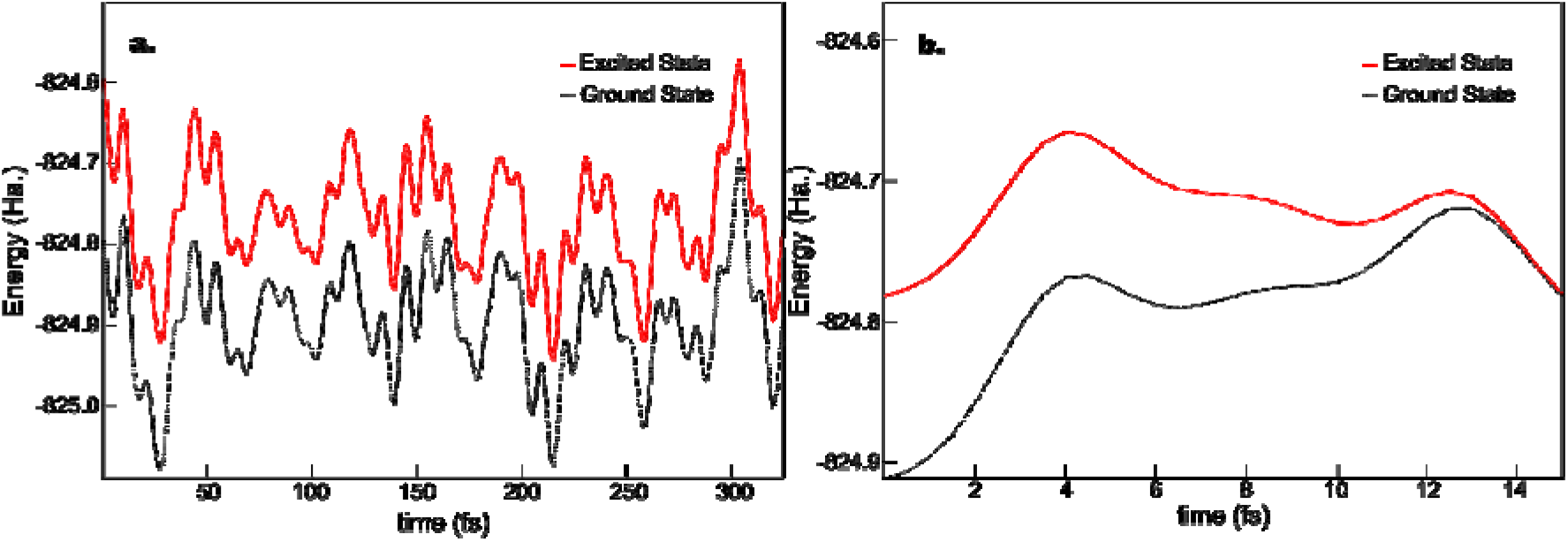
The energy gap between excited and ground state of L-pyro-amm is consistent with the experiments and eventually deactivates via a conical intersection back to the ground state. (a) Shows the energy of the excited and ground state obtained from the excited state molecular dynamics simulations where non-adiabatic decay probabilities are estimated. We see clearly that the energy difference between the excited and ground state is approximately 3 eV (413 nm) consistent with experimental data. (b) Shows the same excited state and ground state energy for another simulation where the system de-activates very quickly within 10s of femtoseconds through a conical intersection where the ground and excited state energies intersect.

## Notes

### Competing Interest Statement

The authors have declared no competing interest.

### Summary of Updates

In the revised manuscript we provide excited state molecular dynamics simulations which directly probe the fluorescence mechanism and further advances this work.

## References

1. A. Shukla, S. Mukherjee, S. Sharma, V. Agrawal, K. V. R. Kishan, P. Guptasarma, A novel UV laser-induced visible blue radiation from protein crystals and aggregates: Scattering artifacts or fluorescence transitions of peptide electrons delocalized through hydrogen bonding? Arch. Biochem. Biophys. 428, 144–153 (2004).

2. G. Rosenman, N. Amdursky, M. Molotskii, D. Aronov, L. Adler-Abramovich, E. Gazit, Blue luminescence based on quantum confinement at peptide nanotubes. Nano Lett. 9, 3111–3115 (2009).

3. F. T. Chan, D. Pinotsi, G. S. K. Schierle, C. F. Kaminski, in Bio-nanoimaging: Protein Misfolding and Aggregation (Elsevier, 2013; http://linkinghub.elsevier.com/retrieve/pii/B9780123944313000134), pp. 147–155.

4. F. T. S. Chan, G. S. Kaminski Schierle, J. R. Kumita, C. W. Bertoncini, C. M. Dobson, C. F. Kaminski, C. V Robinson, C. M. Dobson, J. Beard, P. Das, K. Jansen, M. DeLucia, W.-L. Lin, G. Dolios, R. Wang, C. B. Eckman, D. W. Dickson, M. Hutton, J. Hardy, T. Golde, Protein amyloids develop an intrinsic fluorescence signature during aggregation. Analyst. 138, 2156 (2013).

5. D. Pinotsi, L. Grisanti, P. Mahou, R. Gebauer, C. F. Kaminski, A. Hassanali, G. S. Kaminski Schierle, Proton Transfer and Structure-Specific Fluorescence in Hydrogen Bond-Rich Protein Structures. J. Am. Chem. Soc. 138, 3046–3057 (2016).

6. J. Pansieri, V. Josserand, S. J. Lee, A. Rongier, D. Imbert, M. M. Sallanon, E. Kövari, T. G. Dane, C. Vendrely, O. Chaix-Pluchery, M. Guidetti, J. Vollaire, A. Fertin, Y. Usson, P. Rannou, J. L. Coll, C. Marquette, V. Forge, Ultraviolet–visible–near-infrared optical properties of amyloid fibrils shed light on amyloidogenesis. Nat. Photonics. 13, 473–479 (2019).

7. L. Adler-Abramovich, E. Gazit, The physical properties of supramolecular peptide assemblies: From building block association to technological applications. Chem. Soc. Rev. 43, 7236–7236 (2014).

8. L. Adler-Abramovich, M. Reches, V. L. Sedman, S. Allen, S. J. B. Tendler, E. Gazit, Thermal and chemical stability of diphenylalanine peptide nanotubes: Implications for nanotechnological applications. Langmuir. 22, 1313–1320 (2006).

9. K. Tao, P. Makam, R. Aizen, E. Gazit, Self-assembling peptide semiconductors. Science (80-.). 358 (2017), p. eaam9756.

10. J. M. Andresen, J. Gayán, L. Djoussé, S. Roberts, D. Brocklebank, S. S. Cherny, L. R. Cardon, J. F. Gusella, M. E. Macdonald, R. H. Myers, D. E. Housman, N. S. Wexler, J. Lorimer, J. Porter, F. Gomez, C. Moskowitz, K. P. Gerstenhaber, E. Shackell, K. Marder, G. Penchaszadeh, S. A. Roberts, A. Brickman, D. Brocklebank, J. Gray, S. R. Dlouhy, S. Wiktorski, M. E. Hodes, P. M. Conneally, J. B. Penney, J. F. Gusella, J. H. Cha, M. Irizarry, D. Rosas, S. Hersch, Z. Hollingsworth, A. B. Young, D. E. Housman, M. M. de Young, E. Bonilla, T. Stillings, A. Negrette, S. R. Snodgrass, M. D. Martinez-Jaurrieta, M. A. Ramos-Arroyoh, J. Bickham, J. S. Ramos, F. Marshall, I. Shoulson, G. J. Rey, A. Feigin, N. Arnheim, A. Acevedo-Cruz, L. Acosta, J. Alvir, K. Fischbeck, L. M. Thompson, A. Young, L. Dure, C. J. O’Brien, J. Paulsen, S. P. Moran, D. Krch, P. Hogarth, D. S. Higgins, B. Landwehrmeyer, M. R. Hayden, E. W. Almqvist, R. R. Brinkman, O. Suchowersky, A. Durr, C. C. Dodé, F. Squitieri, P. J. Morrison, M. Nance, C. A. Ross, R. L. Margolis, A. Rosenblatt, G. T. Estrella, D. M. Cabrero, R. J. A. Trent, E. McCusker, A. Novelletto, M. Frontali, J. S. Paulsen, R. Jones, A. Zanko, T. Ashizawa, A. Lazzarini, J. L. Li, V. C. Wheeler, A. L. Russ, G. Xu, J. S. Mysore, T. Gillis, M. Hakky, L. A. Cupples, M. Saint-Hilaire, S. M. Hersch, The relationship between CAG repeat length and age of onset differs for Huntington’s disease patients with juvenile onset or adult onset. Ann. Hum. Genet. 71, 295–301 (2007).

11. D. Pinotsi, A. K. Buell, C. M. Dobson, G. S. Kaminski Schierle, C. F. Kaminski, A Label-Free, Quantitative Assay of Amyloid Fibril Growth Based on Intrinsic Fluorescence. ChemBioChem. 14, 846–850 (2013).

12. T. N. Tikhonova, N. R. Rovnyagina, A. Y. Zherebker, N. N. Sluchanko, A. A. Rubekina, A. S. Orekhov, E. N. Nikolaev, V. V. Fadeev, V. N. Uversky, E. A. Shirshin, Dissection of the deep-blue autofluorescence changes accompanying amyloid fibrillation. Arch. Biochem. Biophys. 651, 13–20 (2018).

13. H. Mori, K. Takio, M. Ogawara, D. J. Selkoe, Mass spectrometry of purified amyloid β protein in Alzheimer’s disease. J. Biol. Chem. 267, 17082–17086 (1992).

14. M. Fändrich, C. M. Dobson, The behaviour of polyamino acids reveals an inverse side chain effect in amyloid structure formation. EMBO J. 21, 5682–90 (2002).

15. R. Dovesi, A. Erba, R. Orlando, C. M. Zicovich-Wilson, B. Civalleri, L. Maschio, M. Rérat, S. Casassa, J. Baima, S. Salustro, B. Kirtman, Quantum-mechanical condensed matter simulations with CRYSTAL. Wiley Interdiscip. Rev. Comput. Mol. Sci. 8, e1360 (2018).

16. P. Giannozzi, O. Andreussi, T. Brumme, O. Bunau, M. Buongiorno Nardelli, M. Calandra, R. Car, C. Cavazzoni, D. Ceresoli, M. Cococcioni, N. Colonna, I. Carnimeo, A. Dal Corso, S. De Gironcoli, P. Delugas, R. A. Distasio, A. Ferretti, A. Floris, G. Fratesi, G. Fugallo, R. Gebauer, U. Gerstmann, F. Giustino, T. Gorni, J. Jia, M. Kawamura, H. Y. Ko, A. Kokalj, E. Kücükbenli, M. Lazzeri, M. Marsili, N. Marzari, F. Mauri, N. L. Nguyen, H. V. Nguyen, A. Otero-De-La-Roza, L. Paulatto, S. Poncé, D. Rocca, R. Sabatini, B. Santra, M. Schlipf, A. P. Seitsonen, A. Smogunov, I. Timrov, T. Thonhauser, P. Umari, N. Vast, X. Wu, S. Baroni, Advanced capabilities for materials modelling with Quantum ESPRESSO. J. Phys. Condens. Matter. 29 (2017), doi:10.1088/1361-648X/aa8f79.

17. M. T. Ruggiero, J. Gooch, J. Zubieta, T. M. Korter, Evaluation of Range-Corrected Density Functionals for the Simulation of Pyridinium-Containing Molecular Crystals. J. Phys. Chem. A. 120, 939–947 (2016).

18. J. Da Chai, M. Head-Gordon, Long-range corrected hybrid density functionals with damped atom-atom dispersion corrections. Phys. Chem. Chem. Phys. 10, 6615–6620 (2008).

19. Y. Noel, C. M. Zicovich-Wilson, B. Civalleri, P. D’Arco, R. Dovesi, Polarization properties of ZnO and BeO: An ab initio study through the Berry phase and Wannier functions approaches. Phys. Rev. B - Condens. Matter Mater. Phys. 65, 1–9 (2002).

20. D. Rocca, R. Gebauer, Y. Saad, S. Baroni, Turbo charging time-dependent density-functional theory with Lanczos chains. J. Chem. Phys. 128 (2008), doi:10.1063/1.2899649.

21. N. Troullier, J. L. Martins, Efficient pseudopotentials for plane-wave calculations. Phys. Rev. B. 43, 1993–2006 (1991).

22. A. D. Becke, Density-functional thermochemistry. III. The role of exact exchange. J. Chem. Phys. 98, 5648–5652 (1993).

23. J. D. Head, M. C. Zerner, A Broyden-Fletcher-Goldfarb-Shanno optimization procedure for molecular geometries. Chem. Phys. Lett. 122, 264–270 (1985).

24. J. VandeVondele, M. Krack, F. Mohamed, M. Parrinello, T. Chassaing, J. Hutter, Quickstep: Fast and accurate density functional calculations using a mixed Gaussian and plane waves approach. Comput. Phys. Commun. 167, 103–128 (2005).

25. S. Goedecker, M. Teter, Separable dual-space Gaussian pseudopotentials. Phys. Rev. B - Condens. Matter Mater. Phys. 54, 1703–1710 (1996).

26. A. D. Becke, Density-functional exchange-energy approximation with correct asymptotic behavior. Phys. Rev. A. 38, 3098–3100 (1988).

27. S. Grimme, J. Antony, S. Ehrlich, H. Krieg, A consistent and accurate *ab initio* parametrization of density functional dispersion correction (DFT-D) for the 94 elements H-Pu. J. Chem. Phys. 132, 154104 (2010).

28. G. Bussi, D. Donadio, M. Parrinello, Canonical sampling through velocity rescaling. J. Chem. Phys. 126 (2007), doi:10.1063/1.2408420.

29. T. Yanai, D. P. Tew, N. C. Handy, A new hybrid exchange-correlation functional using the Coulomb-attenuating method (CAM-B3LYP). Chem. Phys. Lett. 393, 51–57 (2004).

30. J. P. Marcolongo, A. Zeida, J. A. Semelak, N. O. Foglia, U. N. Morzan, D. A. Estrin, M. C. González Lebrero, D. A. Scherlis, Chemical Reactivity and Spectroscopy Explored From QM/MM Molecular Dynamics Simulations Using the LIO Code. Front. Chem. 6, 70 (2018).

31. U. N. Morzan, F. F. Ramírez, M. B. Oviedo, C. G. Sánchez, D. A. Scherlis, M. C. G. Lebrero, Electron dynamics in complex environments with real-time time dependent density functional theory in a QM-MM framework. J. Chem. Phys. 140, 164105 (2014).

32. M. A. Nitsche, M. Ferreria, E. E. Mocskos, M. C. G. Lebrero, GPU accelerated implementation of density functional theory for hybrid QM/MM simulations. J. Chem. Theory Comput. 10, 959–967 (2014).

33. F. Ramírez, · Gonzalo, D. Mirón, M. C. González Lebrero, D. A. Scherlis, QM-MM Ehrenfest dynamics from first principles: photodissociation of diazirine in aqueous solution. Theor. Chem. Acc. 137, 124 (2018).

34. E. Tapavicza, I. Tavernelli, U. Rothlisberger, Trajectory surface hopping within linear response time-dependent density-functional theory. Phys. Rev. Lett. 98, 023001 (2007).

35. J. C. Tully, Molecular dynamics with electronic transitions. J. Chem. Phys. 93, 1061–1071 (1990).

36. M. Barbatti, Nonadiabatic dynamics with trajectory surface hopping method. Wiley Interdiscip. Rev. Comput. Mol. Sci. 1 (2011), pp. 620–633.

37. E. Tapavicza, G. D. Bellchambers, J. C. Vincent, F. Furche, Ab initio non-adiabatic molecular dynamics. Phys. Chem. Chem. Phys. 15 (2013), pp. 18336–18348.

38. C. Adamo, V. Barone, Toward reliable density functional methods without adjustable parameters: The PBE0 model. J. Chem. Phys. 110, 6158–6170 (1999).

39. S. Hirata, M. Head-Gordon, Time-dependent density functional theory within the Tamm-Dancoff approximation. Chem. Phys. Lett. 314, 291–299 (1999).

40. K. Jong, Y. Taghipour Azar, L. Grisanti, A. D. Stephens, S. T. E. Jones, D. Credgington, G. S. Kaminski Schierle, A. A. Hassanali, Low Energy Optical Excitations as an Indicator of Structural Changes Initiated at the Termini of Amyloid Protein. Phys. Chem. Chem. Phys. (2019), doi:10.1039/c9cp04648h.

41. L. Grisanti, D. Pinotsi, R. Gebauer, G. S. Kaminski Schierle, A. A. Hassanali, A computational study on how structure influences the optical properties in model crystal structures of amyloid fibrils. Phys. Chem. Chem. Phys. 19, 4030–4040 (2017).

42. E. P. J. Parrott, J. A. Zeitler, Terahertz time-domain and low-frequency Raman spectroscopy of organic materials. Appl. Spectrosc. 69, 1–25 (2015).

43. Y. K. Law, A. A. Hassanali, Role of Quantum Vibrations on the Structural, Electronic, and Optical Properties of 9-Methylguanine. J. Phys. Chem. A. 119, 10816–10827 (2015).

44. S. Sappati, A. Hassanali, R. Gebauer, P. Ghosh, Nuclear quantum effects in a HIV/cancer inhibitor: The case of ellipticine. J. Chem. Phys. 145 (2016), doi:10.1063/1.4968046.

45. Y. K. Law, A. A. Hassanali, The importance of nuclear quantum effects in spectral line broadening of optical spectra and electrostatic properties in aromatic chromophores. J. Chem. Phys. 148 (2018), doi:10.1063/1.5005056.

46. D. Marx, Erratum: Proton transfer 200 years after von grotthuss: Insights from Ab initio simulations (ChemPhysChem (2006) 7 (1848-1810)). ChemPhysChem. 8, 209–210 (2007).

47. S. Prasad, I. Mandal, S. Singh, A. Paul, B. Mandal, R. Venkatramani, R. Swaminathan, Near UV-Visible electronic absorption originating from charged amino acids in a monomeric protein. Chem. Sci. 8, 5416–5433 (2017).

48. I. Mandal, S. Paul, R. Venkatramani, Optical backbone-sidechain charge transfer transitions in proteins sensitive to secondary structure and modifications. Faraday Discuss. 207, 115–135 (2018).

49. A. Handelman, N. Kuritz, A. Natan, G. Rosenman, Reconstructive Phase Transition in Ultrashort Peptide Nanostructures and Induced Visible Photoluminescence. Langmuir. 32 (2016), pp. 2847–2862.

50. S. K. Joseph, N. Kuritz, E. Yahel, N. Lapshina, G. Rosenman, A. Natan, Proton-Transfer-Induced Fluorescence in Self-Assembled Short Peptides. J. Phys. Chem. A. 123, 1758–1765 (2019).

51. L. L. Del Mercato, P. P. Pompa, G. Maruccio, A. Della Torre, S. Sabella, A. M. Tamburro, R. Cingolani, R. Rinaldi, Charge transport and intrinsic fluorescence in amyloid-like fibrils. Proc Natl Acad Sci U S A. 104, 18019–18024 (2007).

52. K. Jong, L. Grisanti, A. Hassanali, Hydrogen Bond Networks and Hydrophobic Effects in the Amyloid β30-35Chain in Water: A Molecular Dynamics Study. J. Chem. Inf. Model. 57, 1548–1562 (2017).

53. C. Niyangoda, T. Miti, L. Breydo, V. Uversky, M. Muschol, Carbonyl-based blue autofluorescence of proteins and amino acids. PLoS One. 12, e0176983 (2017).

54. A. Stavrinides, E. C. Tatsis, L. Caputi, E. Foureau, C. E. M. Stevenson, D. M. Lawson, V. Courdavault, S. E. O’Connor, Structural investigation of heteroyohimbine alkaloid synthesis reveals active site elements that control stereoselectivity. Nat. Commun. 7 (2016), doi:10.1038/ncomms12116.

55. F. Forneris, D. P. H. M. Heuts, M. Delvecchio, S. Rovida, M. W. Fraaije, A. Mattevi, Structural analysis of the catalytic mechanism and stereoselectivity in Streptomyces coelicolor alditol oxidase. Biochemistry. 47, 978–985 (2008).

56. S. Yamaguchi, H. Kamikubo, K. Kurihara, R. Kuroki, N. Niimura, N. Shimizu, Y. Yamazaki, M. Kataoka, Low-barrier hydrogen bond in photoactive yellow protein. Proc. Natl. Acad. Sci. U. S. A. 106, 440–444 (2009).

57. W. W. Cleland, P. A. Frey, J. A. Gerlt, The low barrier hydrogen bond in enzymatic catalysis. J. Biol. Chem. 273 (1998), pp. 25529–25532.

58. S. Zhou, L. Wang, Unraveling the structural and chemical features of biological short hydrogen bonds. Chem. Sci. 10, 7734–7745 (2019).

